# *In Vivo* Validation of Bimolecular Fluorescence Complementation (BiFC) to Investigate Aggregate Formation in Amyotrophic Lateral Sclerosis (ALS)

**DOI:** 10.1101/2020.10.08.330894

**Authors:** Emily K Don, Alina Maschirow, Rowan A W Radford, Natalie M Scherer, Andres Vidal-Itriago, Alison Hogan, Cindy Maurel, Isabel Formella, Jack J Stoddart, Thomas E Hall, Albert Lee, Bingyang Shi, Nicholas J Cole, Angela S Laird, Andrew P Badrock, Roger S Chung, Marco Morsch

**Affiliations:** Centre for Motor Neuron Disease Research, Faculty of Health and Medical Sciences, Department of Biomedical Science, Macquarie University, Sydney, NSW 2019, Australia; Institute for Molecular Bioscience, The University of Queensland, QLD 4072, Australia

**Keywords:** Bimolecular Fluorescence Complementation, aggregate formation, zebrafish, TDP-43, FUS, ALS

## Abstract

Amyotrophic lateral sclerosis (ALS) is a form of motor neuron disease (MND) that is characterized by the progressive loss of motor neurons within the spinal cord, brainstem and motor cortex. Although ALS clinically manifests as a heterogeneous disease, with varying disease onset and survival, a unifying feature is the presence of ubiquitinated cytoplasmic protein inclusion aggregates containing TDP-43. However, the precise mechanisms linking protein inclusions and aggregation to neuronal loss are currently poorly understood.

Bimolecular Fluorescence Complementation (BiFC) takes advantage the association of fluorophore fragments (non-fluorescent on their own) that are attached to an aggregation prone protein of interest. Interaction of the proteins of interest allows for the fluorescent reporter protein to fold into its native state and emit a fluorescent signal. Here, we combined the power of BiFC with the advantages of the zebrafish system to validate, optimize and visualize of the formation of ALS-linked aggregates in real time in a vertebrate model. We further provide *in vivo* validation of the selectivity of this technique and demonstrate reduced spontaneous self-assembly of the non-fluorescent fragments *in vivo* by introducing a fluorophore mutation. Additionally, we report preliminary findings on the dynamic aggregation of the ALS-linked hallmark proteins Fus and TDP-43 in their corresponding nuclear and cytoplasmic compartments using BiFC.

Overall, our data demonstrates the suitability of this BiFC approach to study and characterize ALS-linked aggregate formation *in vivo*. Importantly, the same principle can be applied in the context of other neurodegenerative diseases and has therefore critical implications to advance our understanding of pathologies that underlie aberrant protein aggregation.

## 1 Introduction

### Aggregate formation in disease

Protein misfolding and aggregation are hallmarks of many different neurodegenerative diseases, however the precise mechanisms linking protein aggregation and neurotoxicity are largely unknown (reviewed in [1],[2],[3]). Importantly, such an aggregation pathology is not exclusively linked to the central nervous system, as many other diseases such as type 2 diabetes, inherited cataracts and myopathies display similar protein abnormalities, making protein aggregation a key concept in biological processes [1],[4].

Protein aggregation is hypothesized to be caused when a particular protein folds into a stable alternative or intermediate conformation and starts to accumulate intra- or extracellularly. Protein misfolding and aggregation is hypothesized to be influenced by several scenarios, including posttranslational modifications like protein hyperphosphorylation or SUMOylation (reviewed in [3],[5]), disease linked-mutations leading to changes in the amino acid sequence (reviewed in [6]), prion-like behaviors (the ability to transmit their misfolded shape onto wild-type variants of the same protein) (reviewed in [4],[7]), pathological increase in protein concentration (also termed “protein supersaturation”) [8], and/or deficiencies in the proteasome or autophagy (protein clearance pathways) [6]. In addition, cellular challenges such as an increase in oxidative stress (reviewed in [9],[10],[11]), increased endoplasmic reticulum stress [12],[13] mitochondrial dysfunction (reviewed in [14],[15]) and alteration of cytoplasmic membrane permeability [16] are all associated with an increase in the severity of the protein aggregation.

### Protein aggregation and neurodegeneration

Amyotrophic lateral sclerosis (ALS) is a neurodegenerative disease, which is characterized by protein aggregates in the upper and lower motor neurons and the degeneration of these neurons (reviewed in [17],[7]). Approximately 10% of ALS patients have a known family history of the disease. To date, gene mutations are the only proven cause of ALS (reviewed in [18]), however the etiology of the disease is complex, with significant clinical heterogeneity and a variable penetrance of ALS-linked mutations. In contrast to the clinical heterogeneity of the disease, mutations in the gene TARDBP (encoding for TDP-43) are unequivocally linked to the development of ALS and TDP-43 inclusions are the key pathological hallmark for nearly all (∼97%) ALS patients [19],[20],[21]. Likewise, mutations in the Fused in Sarcoma (*FUS*) gene account for approximately 4% of familial ALS patients and FUS protein inclusions are also found in a subset of ALS patients (1%) [22],[23],[24],[25]. TDP-43 and FUS are predominately nuclear proteins, which have well characterized nuclear functions, such as transcription, mRNA splicing and poly-adenylation, miRNA biogenesis (reviewed in [26],[27],[28]). Under physiological conditions, both proteins are found in low levels in the cytoplasm [29], where they have emerging roles in mRNA stability and transport, regulation of translation, miRNA processing, stress response, mitochondrial and autophagy regulation and synaptic function (reviewed in [30]). In ALS on the other hand, both TDP-43 and FUS are depleted from the nucleus and mislocalize to the cytoplasm [19],[22], [25]. Clinical verification of TDP-43 and FUS pathology are currently limited to post-mortem tissue examination. However, these histological techniques provide only a static snapshot of the aggregation pattern at predetermined stages of the disease. This significantly limits the ability to investigate the dynamic molecular mechanisms that are believed to trigger aggregate formation, maturation and mislocalization into the cytoplasm.

### A zebrafish model to study neurodegeneration

Zebrafish are an excellent model to study some of the underlying mechanisms as they develop ex-utero (allows for straight-forward genetic modification), are transparent during early age (aiding microscopic analysis) and can be assessed for movement impairment at an early age [31]. For example, zebrafish embryos have previously been used to study the effect of expression of disease-causing mutations in TDP-43, SOD1, FUS, C9orf72 and CCNF [32],[33],[34],[35],[36],[37],[38],[39],[40],[41],[42],[43],[44],[45],[46]. We have previously used the zebrafish system to visualize the intraneuronal localization and spread of human TDP-43 in degenerating motor neurons [47]. However, a limitation of this study was the inability to determine whether the spread of the “pathogenic” TDP-43 released from the dying motor neurons ultimately resulted in interaction with other TDP-43 molecules in neighboring cells, such as the glia or healthy motor neurons. The mechanism through which degeneration spreads, and whether this occurs through prion-like properties, represents a key question in the field.

### Bimolecular Fluorescence Complementation – a tool to visualize protein interactions dynamically

A variety of luminescence-based techniques to trace protein interactions in living cells have been established, including Förster resonance energy transfer (FRET) [48], bioluminescence resonance energy transfer (BRET) [49] and Bimolecular Fluorescence Complementation (BiFC) [50]. BiFC has been previously used to validate protein interactions [51]. It is based on the reassembly of the unfolded, complementary, non-fluorescent N-and C-terminal fragments of a split fluorophore, which are fused to the protein of interest. Upon interaction of the protein of interest, the split fluorophore fragments are brought into spatial proximity (generally <7 nm), enabling the structural complementation of the fluorophore.

BiFC offers unique advantages to study protein-protein interactions such as aggregation, maturation and mislocalization in living cells (reviewed in [51]). The fluorescence complementation results in a specific signal due to the intrinsic fluorescence reconstitution of the non-fluorescent fragments [50]. BiFC does not rely on energy transfer between fluorophores, as seen in FRET and BRET assays, and can be detected by employing conventional fluorescence microscopy [52]. Importantly, the technique can also be adapted to confirm protein interactions within a complex [52]. Potential artifacts, caused by cell lysis or cell fixation, are eliminated, as the protein-protein interactions are studied in living cells [53]. BiFC has previously been used *in vitro* to visualize protein-protein interactions in the early stages of aggregate formation in neurodegenerative disease [54],[55]. BiFC has also been adapted for *in vivo* use in the zebrafish to study protein interactions in cellular pathways during development [56],[57],[58],[59]. Recently, studies have utilized a split red fluorescent protein (RFP) assay in *C*.*elegans* to investigate neuronal synapses [60] and a split GFP complementation assay to visualize alpha-synuclein in zebrafish [61]. In addition, split luciferase and mVenus BiFC assays have been used to visualize α-Synuclein oligomerization in mouse brains [62],[63].

In this study we validated a fluorescence complementation approach to investigate the formation and localization of ALS aggregates *in vivo* by combining BiFC with the advantages of the zebrafish model system. We further provide *in vivo* validation of an optimized BiFC construct and show the selectivity of this approach using competitive injections as a control for non-specific protein interactions, which have previously been reported in BiFC assays [64]. We also demonstrate preliminary results for the dynamic aggregation of the ALS-linked proteins Fus and TDP-43, both wild-type and ALS-linked mutant forms, via fluorescence complementation. Taken together, our results confirm that BiFC can be utilized to study ALS-linked aggregate formation and conceivably spread *in vivo*.

## 2 Material and methods

### 2.1 Zebrafish care

Experiments were conducted under Macquarie University Animal Ethics and Biosafety approvals (2012/050, 2015/034 and 2015/033; 5201401007). Adult wild-type zebrafish (*Danio rerio*) were maintained under standard conditions on a 14:10 light:dark cycle with twice daily feeding of artemia and standard pellet at 28 °C [65]. Larvae were raised in E3 medium (5 mM NaCl, 0.17 mM KCl, 0.33 mM CaCl2 and 0.33 mM MgSO4 buffered to 7.3 pH using Carbonate hardness generator (Aquasonic), no methylene blue) at 28 °C on a 14:10 light:dark cycle.

### 2.2 Generation of constructs

The BiFC fragments chosen for all experiments described here are the amino-terminal (N-terminal) half of the yellow fluorescent protein (YFP) Venus (Venus-1-155, referred to as VN155 hereafter) and the carboxy-terminal (C-terminal) half of the cyan fluorescent protein (CFP) mCerulean (mCerulean-155-239, referred to as CC155 hereafter). These fragments were fused to a protein of interest via a glycine Serine (GGGS)3 flexible linker to allow for the normal biological function of the fusion protein [66]. The constructs were cloned into the mammalian, avian, xenopus and zebrafish expression pCS2+ vector. This vector contains an SP6 promoter allowing for the *in-vitro* transcription of mRNA.

TDP-43 BiFC constructs: Synthesis and subcloning of the gene fragments into pCS2+ plasmids were performed by GenScript (NJ, USA). Wild-type human TDP-43 was N-terminally fused to either VN155 or CC155 via a (GGGS)3 flexible linker. VN155 or CC155 with a (GGGS)3 flexible linker was ordered as controls. Gene fragments were inserted into BamH1 and Xba1 restriction sites of the pCS2+ plasmid. Site directed mutagenesis was performed by GenScript (NJ, USA) to obtain the optimised Venus I152L constructs.

Fus BiFC constructs: VN155 was PCR amplified from mVenus sequence, or CC155 PCR amplified from mCerulean sequence, containing 5’ EcoRI and 3’ EcoRV restriction enzymes sites (see Table S1 for oligonucleotide sequences). Zebrafish Fus was amplified from wild-type embryonic cDNA 24hpf with 5’ EcoRV and 3’ SpeI restriction sites (see Table S1). VN155 or CC155 containing a (GGGS)3 flexible linker was N-terminally fused to Fus in the pCS2+ backbone. Fus was then mutated using a reverse primer encoding the R536G substitution, which serendipitously deleted amino acids 529-532 of the Fus PY-NLS. Herein, this mutation is now referred to as Fus-mutant. Finally, pCS2+ CC155-mKate2 was constructed by excising Fus and subcloning PCR amplified mKate2 containing 5’EcoRV and 3’ SpeI restriction enzyme sites.

**Table 1.**
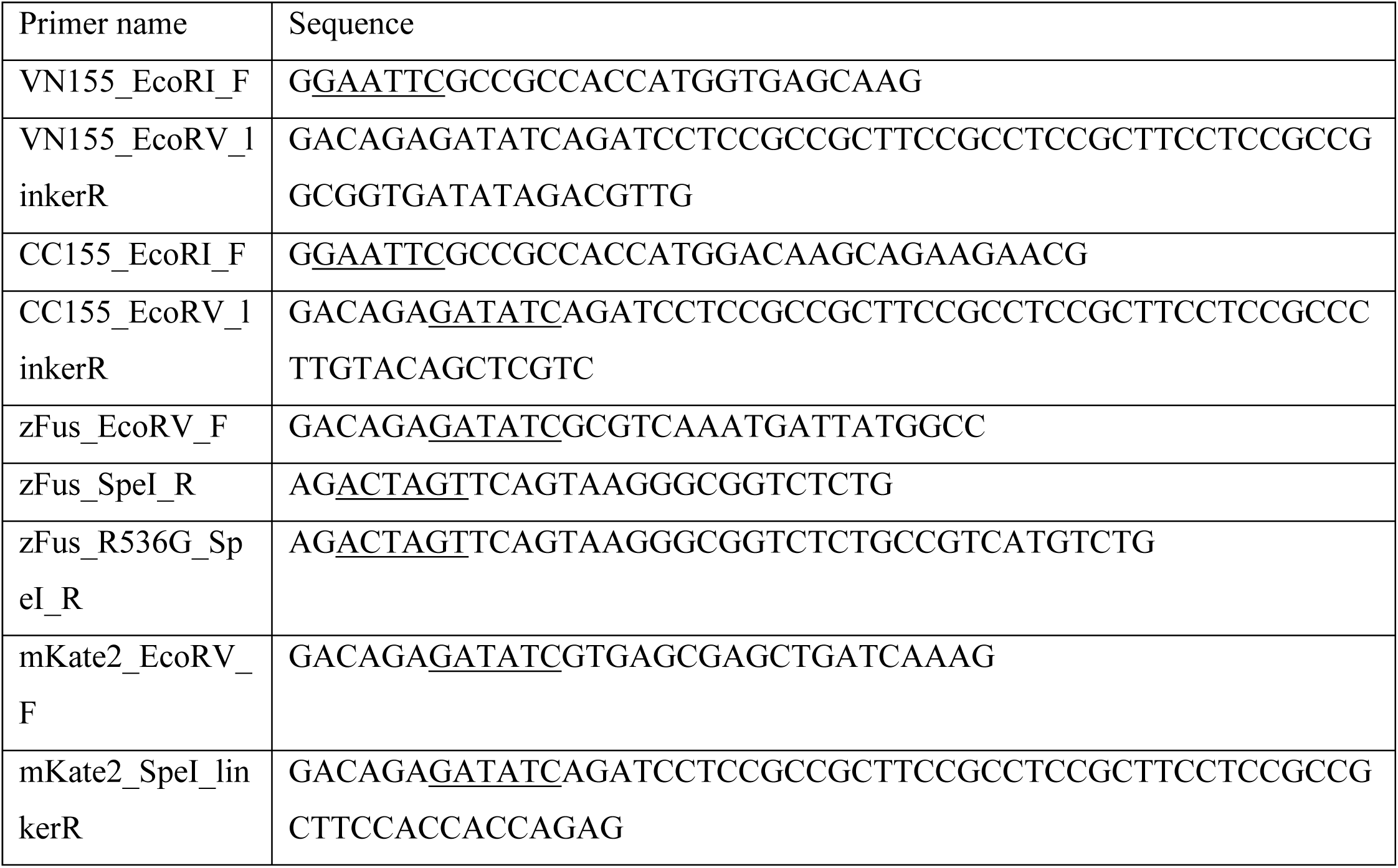
Oligonucleotides used in this study

### 2.3 Generation of mRNA and microinjections

The pCS2+ BiFC constructs were linearized with Not1 and mRNA was transcribed using a mMessage Machine Sp6 Transcription kit (Thermo Fischer Scientific) according to manufactures instructions. Microinjections were performed using a Picospritzer II (Parker Instruments) into the 1-2 cell stage embryo as previously demonstrated [67],[38],[68],[69]. For standard TDP-43 experiments a 1:1 ratio of complementary mRNA (20 pg) was injected with 100 pg H2B-mCerulean3 mRNA (n=583). In control experiments assessing the non-specific fluorescence reconstitution, an adjusted 1:5 ratio of fluorophore fragment control mRNA (CC155) and TDP-43-fused fragment (VN155-TDP-43) was used to adjust for different construct lengths (CC155 and VN155-TDP-43 are encoded by 303 and 1755 base pairs, respectively, three independent experiments, n=36 per experiment). For Fus experiments 100 pg Fus (or 50 pg of CC155-mKate2 in control experiments) and 400 pg H2B-mCerulean3 mRNA was injected (n=3). Images were taken on either a M165FC fluorescent stereomicroscope (Leica) or TCS SP5 Confocal Microscope (Leica) with settings kept constant within experiments.

### 2.4 Plate reader BiFC signal intensity measurement

24 hpf embryos injected with TDP-43 BiFC mRNA were screened for normal morphology and fluorescent TDP-43 expression at a Leica M165 Fluorescence Microscope. Embryos lacking any fluorescent signal were excluded and the remaining embryos which were expressing in similar amounts were placed in a black wall / clear, flat bottom 96-well plate (Corning; 1 fish per well in 200 ml E3). Prior to experiments, embryos were anesthetized by adding tricaine to the wells (MS-222, Sigma-Aldrich, final concentration 0.2 g/L). Intensity of the BiFC signal was measured at 28° C in a PHERAstar FS Microplate Reader (BMG Labtech) using the FI 485 520 optic module, a 7×7 matrix scan and bottom optic mode with a focal height of 3.9 mm. Gain adjustment was performed for independent experiments but kept consistent for multiple plate readings within a single experiment. Autofluorescence of non-injected control embryos was recorded in parallel in the same well plate. For competitive BiFC control experiments, a total of 36 embryos per injection group (standard vs. competitive and wild-type vs. mutant TDP-43, respectively) and 21 control embryos per well plate were measured. For signal/noise ratio experiments, signal and noise BiFC intensities were measured in separate well plates using 36 embryos per injection group (signal VN155 vs. signal VN155-I152L and noise VN155 vs. noise VN155-I152L, respectively) together with 21 non-injected control embryos. All experiments were independently repeated. Gain adjustment was performed based on noise well plate to avoid detection failure and kept consistent for subsequent signal measurement. Data were collected and exported using MARS Data Analysis Software (BMG Labtech). Data was recorded as relative fluorescent units (Supplementary files 1 & 2) and referred to as BiFC intensity in figures.

### 2.5 Confocal microscopy BiFC signal intensity measurement

Tissue-specific imaging at higher resolutions was performed on a Leica SP5 or Leica SP8 confocal microscope following protocols detailed previously [70],[68]. Acquisition was performed with identical gain and laser power settings for each of the treatment groups. The mean fluorescence intensity was measured using ImageJ (freehand selection tool) on maximum projection images acquired in the muscle from three larvae expressing either wild-type or mutant (M337V) TDP-43. Fluorescence intensities were obtained for the whole cell, as well as the cytoplasm and nucleus specifically. To account for any intrinsic differences in fluorescence intensity, the ratio of the cytoplasm and the nucleus for each individual cell was calculated and used for statistical analysis.

### 2.6 Statistical analysis

GraphPad PRISM® (GraphPad Software, Inc.) was used to analyze data and create figures. Statistical analysis on competitive control injection experiments (Figure 1) was performed based on pooled individual data from four independent experiments (n=36) with an unpaired, two-tailed t-test at a 95 % confidence interval. Data was normalized to controls. Statistical analysis on noise and signal experiments (Figure 2) was performed based on pooled individual data from three independent experiments (n=36) with an unpaired, two-tailed t-test at a 95 % confidence interval. Data was normalized to controls. Statistical analysis on the signal to noise ratio was performed on averaged data from three independent experiments with an unpaired, two-tailed t-test and data was normalized to controls. Statistical analysis on the cytoplasmic/nuclear fluorescence intensity signal was performed on pooled data from 3 independent fish with n=37 (wild-type) and n=32 (mutant) cells analyzed with an unpaired, two-tailed t-test at a 95 % confidence interval. Values in figures show mean (bar graphs) or individual data points, and error bars in all figures represent standard deviation (SD), **** = p < 0.0001, *** = p < 0.001, ** = p < 0.01, * = p < 0.05.

**Fig 1.**
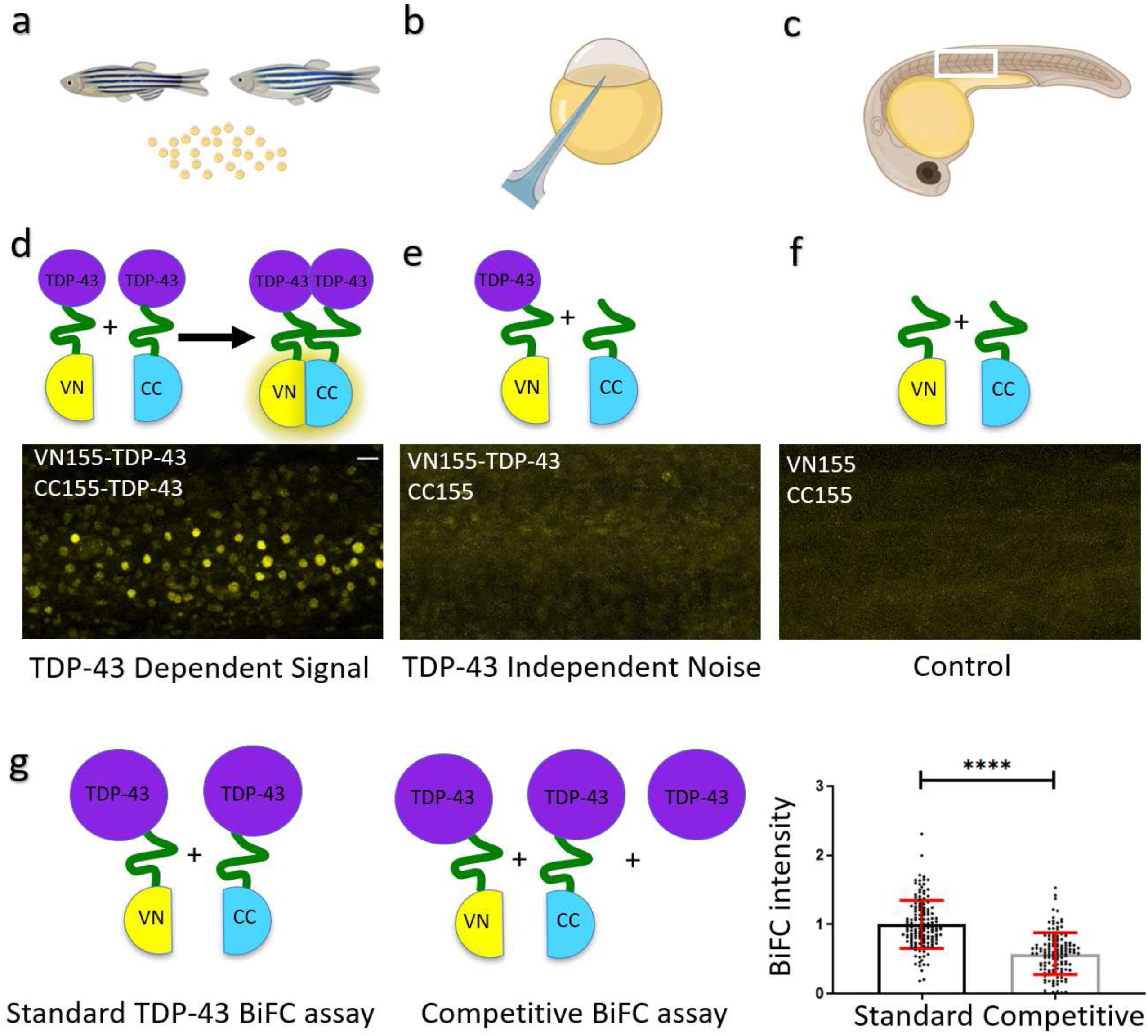
TDP-43 aggregation in zebrafish is specific BiFC assay to determine TDP-43 aggregation. **a-c:** Graphical illustration of the workflow. Male and female zebrafish are set up to collect the fertilized eggs (a). Eggs at the 1-2 cell stage are microinjected (b) with a combination of BiFC mRNA as illustrated below. Embryos are raised and BiFC complementation is visualized using a microscope (c). **d-f:** Top schematics illustrate the different injection combinations. **d:** TDP-43-aggregation and respective fluorescent signal in zebrafish somites at 24hpf. **e:** TDP-43-independent signal (noise). **f:** Control (background) fluorescence at 24 hpf. Scale bar represents 20 µm. **g:** Comparison of BiFC intensity after standard and competitive TDP-43 BiFC injections. The bar diagram shows the quantitative comparison of the normalized BiFC intensity in this fluorescence complementation assay. Dot points represent individual fish and data were pooled from 4 independent experiments. **** = P < 0.0001.

**Fig 2.**
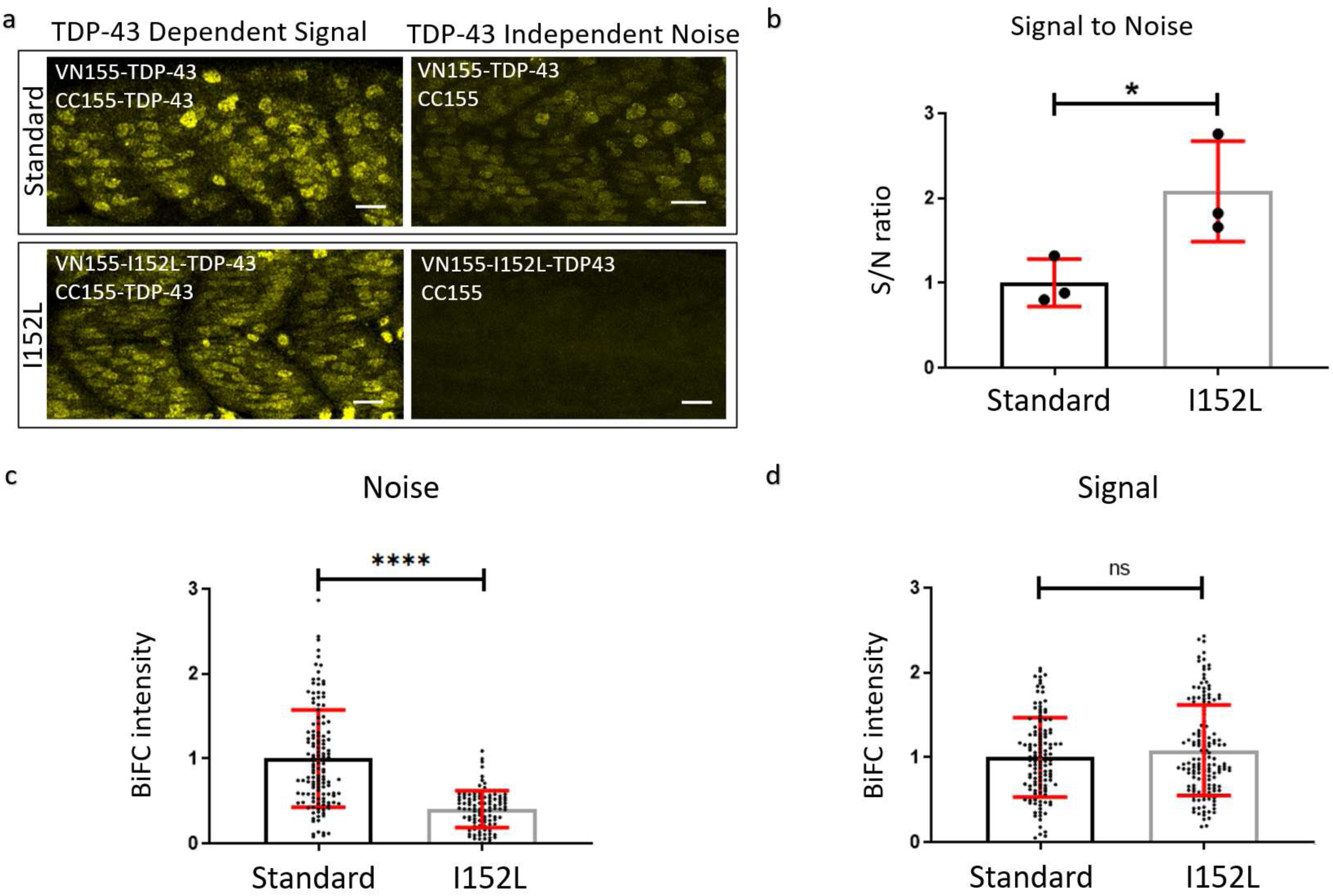
Optimized mVenus fragment (VN155-I152L) increases the signal-to-noise ratio of TDP-43 complementation. **a**: Representative microscope images demonstrating the fluorescence reconstitution in muscle tissue at 24 hpf using VN155-TDP-43 versus VN155-I152L-TDP-43 constructs. Scale bars represent 20 µm. **b**: Quantitative comparison of signal-to-noise ratio using standard VN155 versus VN155-I152L fragments. Data was obtained using an unbiased plate reader. Data pooled and averaged from three independent replicates **c-d:** The background fluorescence (noise) was significantly reduced with the I152L mutation (c), while the maximum fluorescence intensity did not change (d). Data in C-D was obtained using an unbiased plate reader. Data pooled from three independent replicates. * = P < 0.05; **** = P < 0.0001; ns = P > 0.05.

## 3 Results

### BiFC facilitates monitoring of TDP-43 aggregation *in vivo*

In order to determine if BiFC could be used to investigate protein aggregation in zebrafish, the aggregation prone and neurodegenerative disease-linked protein TDP-43 [19] was utilized in our assay. We have previously visualized the full version of the fluorescently tagged human TDP-43 and have determined that wild-type human TDP-43 is predominately located in the nucleus of zebrafish spinal cord motor neurons with approximately 20% found in the cytoplasm [47].

Full-length human wild-type TDP-43 (mRNA) attached with either of the two split-fluorophore halves (CC155 and VN155; Figure 1) was injected into the one-cell stage of the zebrafish eggs to induce expression throughout the embryos. We first performed a dose response curve to establish a dose that did not result in any phenotypical abnormalities such as developmental death or uncharacteristic morphological features in the injected fish. 20 pg was established as an optimal injection dose that did not result in abnormal phenotypes (three independent experiments, minimum n=15 per experiment, Supp Fig 1).

We next assessed if these injections resulted in fluorescence complementation. Clear fluorescence reconstitution was observed in injected embryos 24 hpf (Fig 1d & 3a). Importantly, we could observe a seemingly nuclear localization of the TDP-43 fluorescence signal as we would expect from previous studies with fluorescent full-length TDP-43 [47] (Fig 3c-h). To confirm that the fluorescence signal was specific to TDP-43 interaction, we performed a series of control experiments. Embryos injected with one TDP-43-fused fluorophore fragment and the complementary non-TDP-43-fused fluorophore fragment (VN155-TDP-43 and CC155) resulted in a very dim background expression pattern (TDP-43 independent noise, Fig 1e). Likewise, embryos injected with fluorophore fragments not fused to TDP-43 (VN155 and CC155) displayed negligible fluorescence, which was distributed homogeneously and clearly distinguishable from the signal resulting from fluorophore fragments fused to TDP-43 (Fig 1f). These control experiments demonstrated that spontaneous, unspecific complementation of fluorophore fragments (noise) was negligible whilst TDP-43-fused BiFC fragments showed a clear propensity for fluorescence reconstitution, likely indicating enhanced aggregate propensities for TDP-43 and the specificity of the assay.

**Fig 3.**
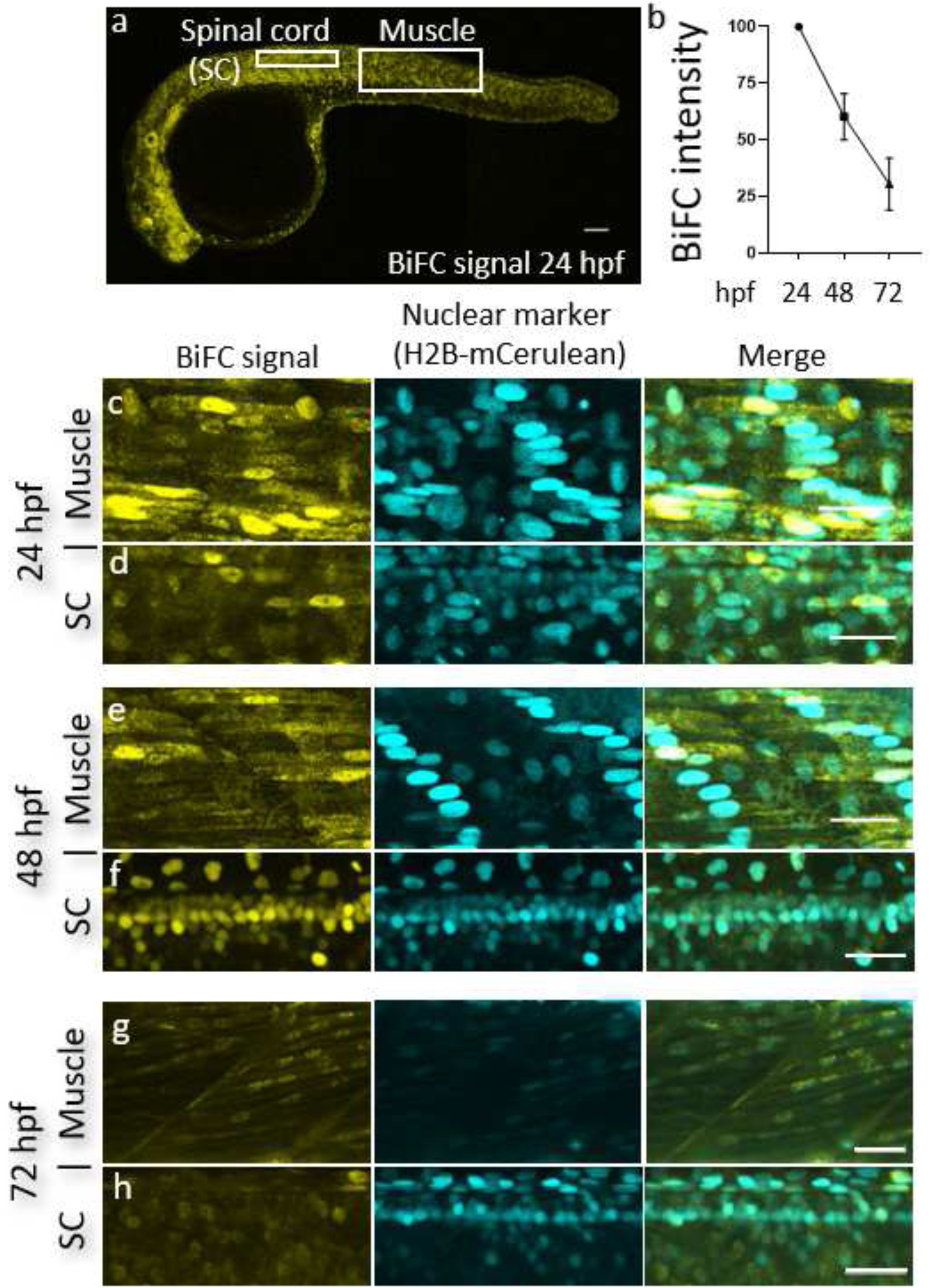
TDP-43 BiFC can be detected in muscle cells and motor neurons and is predominantly nuclear. **a**: A representative image of a 24 hpf embryo injected with our TDP-43 BiFC constructs. Scale bar: 200 µm **b:** Fluorescent intensity as a percentage of signal at 24 hpf. **c-d**: Representative pictures of the human wild-type TDP-43 BiFC signal at 24 hpf in muscle cells (c) and spinal cord motor neurons (d). **e-f**: Representative pictures of the human wild-type TDP-43 BiFC signal at 48 hpf in muscle cells (e) and spinal cord motor neurons (f). **g-h:** Representative pictures of the human wild-type TDP-43 BiFC signal at 72 hpf in muscle cells (g) and spinal cord motor neurons (h). Scale bar: 30 µm.

If the fluorescence reconstitution is a result of aggregating TDP-43 in our *in vivo* system, we would expect that additional supplementation with TDP-43 mRNA not coupled to a fluorophore half results in reduced BiFC intensity. To test this, non-fluorescent TDP-43 was injected with or without the same amount of the BiFC-halves (Fig 1g). Here, we applied a microplate reader-based approach to measure signal intensities in an unbiased fashion. Injection of 10 pg TDP-43 control mRNA (non-fused) in addition to the standard BiFC halves (dose of 10 pg VN155-TDP-43 + 10 pg CC155-TDP-43 mRNA) resulted in a decreased BiFC intensity as compared to standard injections (10 pg VN155-TDP-43 + 10 pg CC155-TDP-43; Fig 1d). The BiFC intensity in these injections was significantly dimmer (0.58 units ± 0.30, three independent experiments, n=36 per experiment) compared to the signal in standard-TDP-43-BiFC-injected embryos (1 unit ± 0.35, three independent experiments, n=36 per experiment; TDP-43 fused to fluorophore-fragments only; absolute intensity divided by auto-fluorescence intensity in non-injected controls and normalized to standard) (Fig 1g).

### Fluorescence optimization *in vivo* using an isoleucine substitution

To further optimize the TDP-43 BiFC efficiency and reduce background (non-specific) fluorescence, a reported optimization for mVenus-based BiFC systems was exploited [64]. This optimization involves an I152L substitution in the N-terminal mVenus fragment which reduces the self-assembly of the fluorophore and should therefore reduce the background non-specific fluorescence of the assay [64]. To determine if the approach would work for our *in vivo* TDP-43 BiFC system, the mVenus I152L substitution was introduced into our constructs. To compare the levels of TDP-43-independent self-assembly of the fluorophore, mRNA injections of TDP-43 fused to either the standard (VN155-TDP-43) or the optimized (VN155-I152L-TDP-43) mVenus fragment were performed in combination with the mCerulean (CC155) control (not fused to TDP-43; Fig 2a, right column). In parallel, the intensity of the TDP-43-dependent fluorescence signal was assessed by injecting TDP-43 fused either to standard (VN155-TDP-43) or optimized (VN155-I152L-TDP-43) mVenus fragments together with the complementary, TDP-43-fused mCerulean (CC155-TDP-43) fragment (Fig 2a, left column). An unbiased plate reader approach was used to determine fluorescence intensities. The optimized I152L substitution decreased non-specific background fluorescence (TDP-43 independent noise) and resulted in a 2-fold increased signal-to-noise ratio as compared to the standard VN155-TDP-43 construct (average mean from three pooled independent experiments, normalized to standard: 2.08 ± 0.59 and 0.99 ± 0.29, respectively, three independent experiments, n=36 per experiment) without diminishing the signal intensity (Fig 2b-d). In summary, the data demonstrate that fluorescence reconstitution in our mRNA-based TDP-43 BiFC assay is TDP-43-interaction-specific, and that the introduction of the I152L substitution further improves the signal to noise ratio 2-fold by reducing the non-specific fluorescence reconstitution.

### TDP-43 BiFC accumulation is tissue and compartment specific

Since mRNA levels are highly regulated during zebrafish early development [71], the BiFC system employed here provides a transient BiFC signal, as the injected constructs are degraded progressively. To determine the duration of TDP-43 BiFC expression, embryos were microinjected at one-cell stage and raised in the dark incubator. The brightness of the BiFC signal was assessed at multiple time points during that period, and individual fish were imaged at consecutive days using identical acquisition parameters. As expected, the fluorescence intensity peaked at 24-30 hpf and faded gradually during the following days (Fig 3b). At 72 hpf the BiFC intensity declined to ∼30 % of the 24 hpf signal, indicating the expected mRNA degradation and ensuing fluorescent reduction (n=5, Fig 3b, c-h). This signal loss is in line with the expected growth-dependent decline in the levels of cellular mRNA and proteins in live tissue, particularly in early developmental stages [72],[71].

TDP-43 has been previously reported to localize primarily in the nucleus [73],[47]. To determine the localization of our TDP-43 BiFC constructs more precisely, co-injections were performed with mRNA that labels the nucleus of the cells (human nuclear protein Histone H2B fused to full length mCerulean3; H2B-mCerulean3). At 24, 48- and 72-hours post fertilization (hpf) the BiFC signal could be clearly observed in the muscle tissue as well as spinal cord motor neurons (n=583, Fig 3a & c-h). Higher magnification confocal microscopy revealed that exogenous wild-type TDP-43 was predominantly located in the nucleus, whilst lower levels of the protein were also present in the cytoplasm, resembling the expression pattern of endogenous TDP-43 ([73], Fig 3c-h). To further analyze the compartmentalization and (mis)localization of TDP-43 we next assessed the BiFC fluorescence expression of wild-type versus mutant (M337V) TDP-43 (Fig 4c-f). Quantitative analysis of confocal z-projections revealed that the mutant form of TDP-43 had a higher propensity to accumulate in the cytoplasm compared to wild-type TDP-43 (Fig 4b). This shift in the cytoplasmic to nuclear fluorescence intensity was independent of the overall fluorescence expression in our fish (Fig 4b).

**Fig 4.**
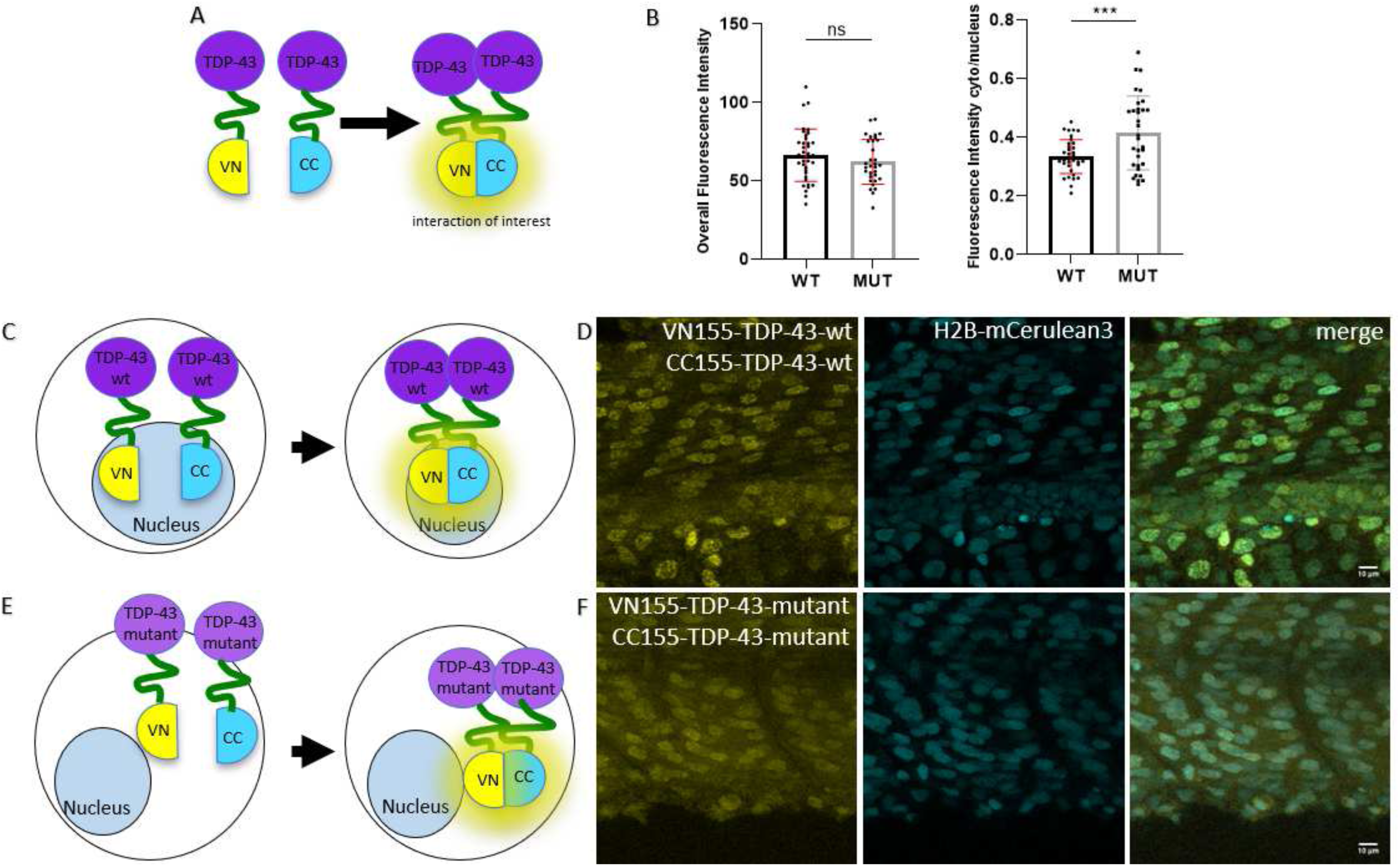
Mutant TDP-43 BiFC is mislocalized to the cytoplasm. **a, c, f:** Schemes of fluorophores used in this TDP-43 BiFC assays. b,**c:** Quantification of fluorescence intensity demonstrates that mutant TDP-43 BiFC is shifted to the cytoplasm with no change to overall fluorescence intensity. **f**: Representative fluorescence images of the wild-type and mutant TDP-43 BiFC signal at 36 hpf in the somites over the yolk extension. Scale bars represent 10 µm.

### Characterization of the interactions of mutant and wild-type Fus using BiFC

To assess if the fluorescence complementation approach is relevant for other ALS proteins, we next tested the ALS-associated protein Fus (Fig 5a). Wild-type Fus BiFC mRNA constructs were injected into the one-cell stage of zebrafish embryos. Injections of VN155-Fus and CC155-Fus resulted, similar to the TDP-43 assay, in successful fluorescence complementation, demonstrating that wild-type Fus proteins interact and accumulate as distinct fluorescent puncta throughout the muscle cells (Figure 5b & c).

**Fig 5.**
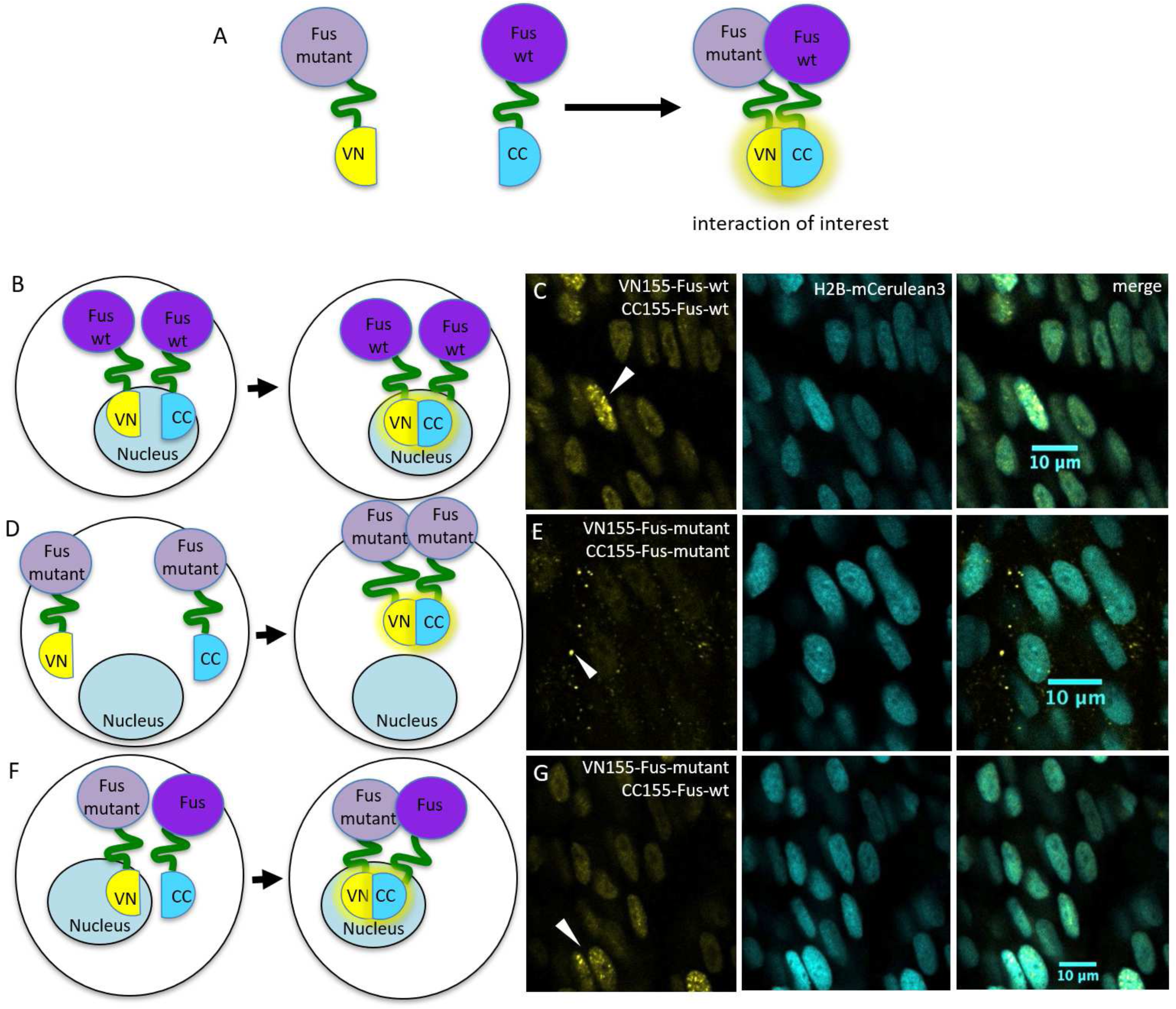
Wild-type and cytoplasmic localized mutant Fus BiFC assay in zebrafish. **a, b, d, f:** Schemes of fluorophores used in this Fus BiFC assays. **c**: Representative fluorescence images of the zebrafish wild-type Fus BiFC signal at 28 hpf in the somites over the yolk extension. **e**: Mutant Fus BiFC signal is observed as discrete puncta in the cytoplasm. **g**: The combination of mutant and wild-type Fus constructs reconstitutes primarily nuclear. Scale bars represent 10 µm.

Disease-linked mutations in *FUS* have been shown to lead to mislocalization of the predominately nuclear FUS protein into the cytoplasm and are a potential cause of pathological mechanism in ALS [22]. Therefore, we next generated an endogenous zebrafish *fus* construct that contained an equivalent human pathological FUS mutation (human FUS^R521G^ aligns with zebrafish Fus^R536G^) and a non-functional nuclear localisation signal (to achieve Fus protein (mis)localization into the cytoplasm). Indeed, injections of the mutant FUS constructs VN155-Fus and CC155-Fus resulted in fluorescence complementation in the cytoplasm as discrete puncta/aggregates (n=3, Fig 5d & e; note the lack of nuclear BiFC signal compared to wild-type Fus). When we next co-expressed wild-type Fus and mutant Fus together, we found that these two proteins interact albeit being determined for different cellular compartments (n=3, wild-type nuclear and mutant cytoplasmic; Fig 5f & g). Additionally, we observed that fluorescence complementation was predominantly restricted to the nucleus, indicating that wild-type Fus has not only the capacity to interact with mutant Fus but conceivably recruits mutant Fus into the nucleus (Fig 5f & g).

To verify that these interactions are specific to Fus-Fus protein interactions and not due to non-specific complementation of the fluorophore fragments, wild-type and mutant Fus N-terminal fluorophore fragments (VN155-Fus) were co-injected with the C-terminal mCerulean fluorophore fragment fused to an mKate2 fluorophore (CC155-mKate2; red fluorescence). This approach let us observe the red fluorescence independent of complementation but would result in yellow mVenus complementation if the interaction of the fluorophore halves was non-specific. Indeed, red fluorescence was visualized throughout the embryo, indicating successful injection and maturation of the fluorophore (Supp Fig 2). Importantly, little to no background fluorescence complementation was observed and no yellow fluorescence reconstitution was detected (Supp Fig 2), highlighting that the Fus fluorescence complementation was due to the specific Fus-Fus protein interactions. Taken together, these results demonstrate that BiFC can be applied to different ALS-linked proteins and effectively used to investigate their aggregate characteristics *in vivo*.

## 4 Discussion

We have optimized and verified a novel BiFC assay for the visualization of ALS-linked protein aggregation in zebrafish. Our BiFC approach provides confirmation that the two neurodegenerative disease-linked proteins, TDP-43 and Fus, undergo fluorescence complementation *in vivo*. Additionally, our data highlights that the BiFC signal could be significantly improved using fluorophore optimization and overall demonstrates the successful implementation of BiFC to observe mislocalization and aggregate formation.

Previous studies have utilized the BiFC system in cell culture for visualization of neurodegenerative disease-linked proteins and to find unprecedented insights into the aggregation propensities of disease aggregates. Tau aggregation and interactions have been studied quantitatively in cell culture using a split GFP/BiFC based approach [54],[74],[55]. This approach revealed that tau aggregation is favored when tau is phosphorylated at S396/S404 in the presence of increased GSK3beta activity [54]. A similar approach has identified that specific amino-acids in the head and tail portions of the N-terminal domain of TDP-43 (E17A; E21A; R52A; R55A) mediate the interactions necessary for its self-assembly [75] and that the self-assembly of FUS is mediated through its low-complexity domain (amino acids 1-214) [76],[77]. BiFC approaches have also been adapted to *in vivo* systems to study morphogen gradients and protein interactions in developmental biology pathways [56],[57],[58],[59]. More recently, BiFC has been implemented to observed neurological processes [60],[61]. In *C*.*elegans*, a split RFP assay differentially labeled subsets of synapses between the same interneurons and different neuronal partners, and demonstrated the *clr-1* gene is required for the formation of both sets of synapses [60]. Vicario et al., also demonstrated that Parkinson’s disease related mutations in α-synuclein resulted in altered sub-mitochondrial localization of the protein using a BiFC approach [61]. These studies highlight the suitability and value of split-fluorescence approaches to uncover some of the fundamental molecular mechanisms that drive protein accumulation.

To verify if BiFC can be used to study the process of aggregate formation in zebrafish, we expressed both TDP-43 and Fus BiFC fusion proteins transiently in zebrafish embryos. Our study revealed fluorescence complementation of wild-type TDP-43 and wild-type Fus and the development of intra-nuclear aggregates, whilst mutant Fus BiFC fusion proteins demonstrated fluorescence complementation in cytoplasmic aggregates as previously reported in cell culture BiFC experiments [75],[77]. Interestingly, co-expression of wild-type and mutant Fus BiFC fusion proteins resulted in fluorescence complementation in intra-nuclear aggregates, not cytoplasmic aggregates, contrary to previously reported findings from cell culture studies and post-mortem tissue [77],[78]. However, the Matsumoto et al. cell culture study did not use a BiFC approach and one possibility is that the intra-nuclear aggregation could represent an early phenomenon (nuclear maturation of the protein followed by cytoplasmic shuttling) that is too late to observe in post-mortem tissue or cell culture models. These preliminary differences highlight the importance of studying these dynamic processes of aggregate formation and maturation longitudinally in living cells/organism such as the zebrafish. Our *in vivo* analysis of TDP-43 compartmentalization also confirmed previous cell culture observations that described increased mislocalization of mutant forms of TDP-43 [79]. As expected, the M337V mutation induced cytoplasmic mislocalization of TDP-43 in our zebrafish (Fig 4). This further demonstrates the suitability of our BiFC approach to study protein interactions and mislocalization in a living animal.

Some limitations of the zebrafish based BiFC system described in this study are the transient nature and ubiquitous expression that result from mRNA injections. During zebrafish early development mRNA levels are highly regulated and the injected constructs, which encode the BiFC proteins, are degraded quickly [71]. Therefore, the applied mRNA system, which has the advantage of rapid detection, is transient and only allows for the detection of fluorescence in the very early stages of development (Fig 3). DNA constructs designed to induce stable transgenesis are needed to allow for the longer-term visualization of disease aggregate maturation and its potential mislocalization. Additionally, in order to study the longer-term effects of ALS aggregates and their precise contribution to neurodegeneration the BiFC constructs could be expressed in neurons and glia respectively. Whilst we tested the toxicity of the BiFC constructs in our assay (Supp Fig 1), the general toxicity level is likely related to the protein of interest (TDP-43 in our assay). Nevertheless, we employed a series of assays to test for the specificity of these interactions and demonstrate that TDP-43 and Fus complementation occurs only when the proteins are fused to the non-fluorescent halves, and that this signal could be increased 2-fold by introducing a previously reported mVenus mutation (I152L) [64] (Fig 2). These results demonstrate that the fluorescence complementation is both a specific and reliable tool to detect ALS-linked protein aggregation *in vivo* in zebrafish. Notably this approach is not limited to ALS aggregates and may present a unique opportunity to investigate propagation of aggregates in other neurodegenerative diseases (such as α-synuclein in Parkinson’s disease; [80]).

The specificity and non-reversible nature of the fluorescence complementation offer unique opportunities to study aggregate maturation and the potential spread of TDP-43 *in vivo*. Current imaging techniques are limited by the quenching of an existing fluorescence signal after neurodegeneration (i.e. due to uptake by microglia; [47]). The BiFC approach may open novel avenues to test the hypothesis that TDP-43 can spread horizontally throughout the spinal cord. Our zebrafish BiFC assay could also be adapted to study stress-granule formation in real time *in vivo*. Stressors involved in disease onset and progression (such as oxidative stress) could be applied to the embryos to determine if and how physiologically relevant cellular stress can exacerbate protein aggregation. Furthermore, a current hypothesis in the field is that TDP-43 aggregates form frequently through a process called liquid-liquid-phase-separation (LLPS; [81]). This process is reversible and allows proteins to form membrane-less sub-compartments where specific short-term actions such as RNA processing can be performed rapidly. However, it is now believed that under certain conditions these liquid droplets can form into tightly clustered aggregates that can become pathological [82]. To the best of our knowledge, this BiFC approach may be one of the very few ways to assess how recruitment of mutant or pathological forms of TDP-43 can directly influence the droplet behaviour of wild-type TDP-43 (or vice versa). Particularly transparent zebrafish offer some unique advantages to examine these pathways in detail as zebrafish embryos can be generated in large numbers, allow high-resolution fluorescence microscopy, and ultimately can provide a useful medium-throughput preclinical platform to screen for molecules that can modulate the aggregation of disease relevant proteins in the future.

## 5 Conclusions

Protein aggregation is hypothesized to underlie a variety of degenerative human diseases. However, the mechanistic connection between the processes of protein aggregation, tissue degeneration and the progression and spread of degeneration is not well understood. Pathological aggregates likely vary across a structural spectrum ranging from small unstructured oligomers to well defined cross-beta-sheet amyloid fibrils. Which forms of aggregate are most toxic, and the mechanisms underlying this proteotoxicity are of immense interest to the field. Identification of the mechanistic and molecular underpinnings, and identifying small molecules with the capacity to modify these processes, may be an important step towards a potential therapeutic intervention applicable to ALS and a range of other human proteinopathies.

## Declarations

### Ethics approval

Experiments were conducted under Macquarie University Animal Ethics and Biosafety approvals (2012/050, 2015/034 and 2015/033; 5201401007).

### Availability of data and material

Plasmids are available via AddGene.

### Authors’ contributions

APB, AM, ED, CM and MM performed the experiments and created the figures, ED, APB and MM wrote the manuscript, all other authors edited the text, performed literature search and provided valuable comments.

## Abbreviations

ALS: Amyotrophic Lateral Sclerosis
BiFC: Bimolecular Fluorescence Complementation
BRET: Bioluminescence Resonance Energy Transfer
CFP: Cyan Fluorescent Protein
FRET: Förster Resonance Energy Transfer
GFP: Green Fluorescence Protein
MND: Hours post fertilization hpf Motor Neuron Disease
RFP: Red Fluorescent Protein
YFP: Yellow Fluorescent Protein

## Acknowledgements

We wish to thank the Snow Foundation for their generous support towards establishing the transgenic zebrafish facility at Macquarie University and continued support of the researchers. We also wish to thank the zebrafish facility staff (past and present) for assistance in zebrafish care.

## References

1. Carrell RW, Lomas DA (1997) Conformational disease. The Lancet 350 (9071):134–138

2. Soto C (2003) Unfolding the role of protein misfolding in neurodegenerative diseases. Nature Reviews Neuroscience 4 (1):49–60. doi:10.1038/nrn1007

3. Tenreiro S, Eckermann K, Outeiro TF (2014) Protein phosphorylation in neurodegeneration: friend or foe? Frontiers in molecular neuroscience 7:42

4. Frost B, Diamond MI (2010) Prion-like mechanisms in neurodegenerative diseases. Nature Reviews Neuroscience 11 (3):155

5. Maurel C, Dangoumau A, Marouillat S, Brulard C, Chami A, Hergesheimer R, Corcia P, Blasco H, Andres C, Vourc’h PJMn (2018) Causative genes in amyotrophic lateral sclerosis and protein degradation pathways: a link to neurodegeneration. 55 (8):6480–6499

6. Blokhuis AM, Groen EJN, Koppers M, van den Berg LH, Pasterkamp RJ (2013) Protein aggregation in amyotrophic lateral sclerosis. Acta neuropathologica 125 (6):777–794. doi:10.1007/s00401-013-1125-6

7. McAlary L, Plotkin SS, Yerbury JJ, Cashman N (2019) Prion-Like Propagation of Protein Misfolding and Aggregation in Amyotrophic Lateral Sclerosis. Frontiers in molecular neuroscience 12:262

8. Yerbury JJ, Ooi L, Blair IP, Ciryam P, Dobson CM, Vendruscolo MJNl (2019) The metastability of the proteome of spinal motor neurons underlies their selective vulnerability in ALS. 704:89–94

9. Lin MT, Beal MF (2006) Mitochondrial dysfunction and oxidative stress in neurodegenerative diseases. Nature 443 (7113):787–795. doi:10.1038/nature05292

10. Barber SC, Shaw PJ (2010) Oxidative stress in ALS: Key role in motor neuron injury and therapeutic target. Free Radical Biology and Medicine 48 (5):629–641. doi:https://doi.org/10.1016/j.freeradbiomed.2009.11.018

11. Lévy E, El Banna N, Baïlle D, Heneman-Masurel A, Truchet S, Rezaei H, Huang M-E, Béringue V, Martin D, Vernis L (2019) Causative Links between Protein Aggregation and Oxidative Stress: A Review. Int J Mol Sci 20 (16):3896. doi:10.3390/ijms20163896

12. Atkin JD, Farg MA, Walker AK, McLean C, Tomas D, Horne MK (2008) Endoplasmic reticulum stress and induction of the unfolded protein response in human sporadic amyotrophic lateral sclerosis. Neurobiology of Disease 30 (3):400–407. doi:https://doi.org/10.1016/j.nbd.2008.02.009

13. Medinas DB, Rozas P, Martínez Traub F, Woehlbier U, Brown RH, Bosco DA, Hetz C (2018) Endoplasmic reticulum stress leads to accumulation of wild-type SOD1 aggregates associated with sporadic amyotrophic lateral sclerosis. Proceedings of the National Academy of Sciences 115 (32):8209–8214. doi:10.1073/pnas.1801109115

14. Sasaki S, Iwata M (1996) Dendritic synapses of anterior horn neurons in amyotrophic lateral sclerosis: an ultrastructural study. Acta neuropathologica 91 (3):278–283

15. Muyderman H, Chen T (2014) Mitochondrial dysfunction in amyotrophic lateral sclerosis - a valid pharmacological target? Br J Pharmacol 171 (8):2191–2205. doi:10.1111/bph.12476

16. Drolle E, Negoda A, Hammond K, Pavlov E, Leonenko Z (2017) Changes in lipid membranes may trigger amyloid toxicity in Alzheimer’s disease. PLoS One 12 (8):e0182194

17. Wood J, Beaujeux T, Shaw P (2003) Protein aggregation in motor neurone disorders. Neuropathology and applied neurobiology 29 (6):529–545

18. Alsultan AA, Waller R, Heath PR, Kirby J (2016) The genetics of amyotrophic lateral sclerosis: current insights. Degenerative neurological and neuromuscular disease 6:49

19. Neumann M, Sampathu DM, Kwong LK, Truax AC, Micsenyi MC, Chou TT, Bruce J, Schuck T, Grossman M, Clark CM (2006) Ubiquitinated TDP-43 in frontotemporal lobar degeneration and amyotrophic lateral sclerosis. Science 314 (5796):130–133

20. Arai T, Hasegawa M, Akiyama H, Ikeda K, Nonaka T, Mori H, Mann D, Tsuchiya K, Yoshida M, Hashizume Y (2006) TDP-43 is a component of ubiquitin-positive tau-negative inclusions in frontotemporal lobar degeneration and amyotrophic lateral sclerosis. Biochemical and biophysical research communications 351 (3):602–611

21. Sreedharan J, Blair IP, Tripathi VB, Hu X, Vance C, Rogelj B, Ackerley S, Durnall JC, Williams KL, Buratti EJS (2008) TDP-43 mutations in familial and sporadic amyotrophic lateral sclerosis. 319 (5870):1668–1672

22. Neumann M, Rademakers R, Roeber S, Baker M, Kretzschmar HA, Mackenzie IR (2009) A new subtype of frontotemporal lobar degeneration with FUS pathology. Brain 132 (11):2922–2931

23. Vance C, Rogelj B, Hortobágyi T, De Vos KJ, Nishimura AL, Sreedharan J, Hu X, Smith B, Ruddy D, Wright P (2009) Mutations in FUS, an RNA processing protein, cause familial amyotrophic lateral sclerosis type 6. Science 323 (5918):1208–1211

24. Kwiatkowski TJ, Bosco D, Leclerc A, Tamrazian E, Vanderburg C, Russ C, Davis A, Gilchrist J, Kasarskis E, Munsat T (2009) Mutations in the FUS/TLS gene on chromosome 16 cause familial amyotrophic lateral sclerosis. Science 323 (5918):1205–1208

25. Radford RA, Morsch M, Rayner SL, Cole NJ, Pountney DL, Chung RS (2015) The established and emerging roles of astrocytes and microglia in amyotrophic lateral sclerosis and frontotemporal dementia. Frontiers in cellular neuroscience 9 (414). doi:10.3389/fncel.2015.00414

26. Ederle H, Dormann D (2017) TDP-43 and FUS en route from the nucleus to the cytoplasm. FEBS letters 591 (11):1489–1507

27. Shang Y, Huang EJ (2016) Mechanisms of FUS mutations in familial amyotrophic lateral sclerosis. Brain research 1647:65–78

28. Ratti A, Buratti E (2016) Physiological functions and pathobiology of TDP-43 and FUS/TLS proteins. Journal of neurochemistry 138:95–111

29. Kapeli K, Martinez FJ, Yeo GWJHg (2017) Genetic mutations in RNA-binding proteins and their roles in ALS. 136 (9):1193–1214

30. Birsa N, Bentham MP, Fratta P Cytoplasmic functions of TDP-43 and FUS and their role in ALS. In: Seminars in cell & developmental biology, 2019. Elsevier,

31. Kimmel CB, Ballard WW, Kimmel SR, Ullmann B, Schilling TF (1995) Stages of embryonic development of the zebrafish. Developmental dynamics 203 (3):253–310

32. Armstrong GAB, Liao M, You Z, Lissouba A, Chen BE, Drapeau P (2016) Homology directed knockin of point mutations in the zebrafish tardbp and fus genes in ALS using the CRISPR/Cas9 system. PLoS One 11 (3)

33. Armstrong GA, Drapeau P (2013) Loss and gain of FUS function impair neuromuscular synaptic transmission in a genetic model of ALS. Human molecular genetics 22 (21):4282–4292

34. Asakawa K, Handa H, Kawakami KJNc (2020) Optogenetic modulation of TDP-43 oligomerization accelerates ALS-related pathologies in the spinal motor neurons. 11 (1):1–16

35. Bosco DA, Lemay N, Ko HK, Zhou H, Burke C, Kwiatkowski Jr TJ, Sapp P, McKenna-Yasek D, Brown Jr RH, Hayward LJ (2010) Mutant FUS proteins that cause amyotrophic lateral sclerosis incorporate into stress granules. Human molecular genetics 19 (21):4160–4175

36. Ciura S, Lattante S, Le Ber I, Latouche M, Tostivint H, Brice A, Kabashi E (2013) Loss of function of C9orf72 causes motor deficits in a zebrafish model of amyotrophic lateral sclerosis. Annals of neurology 74 (2):180–187

37. Hewamadduma CA, Grierson AJ, Ma TP, Pan L, Moens CB, Ingham PW, Ramesh T, Shaw PJ (2013) Tardbpl splicing rescues motor neuron and axonal development in a mutant tardbp zebrafish. Human molecular genetics 22 (12):2376–2386

38. Hogan AL, Don EK, Rayner SL, Lee A, Laird AS, Watchon M, Winnick C, Tarr IS, Morsch M, Fifita JA (2017) Expression of ALS/FTD-linked mutant CCNF in zebrafish leads to increased cell death in the spinal cord and an aberrant motor phenotype. Human molecular genetics 26 (14):2616–2626

39. Kabashi E, Lin L, Tradewell ML, Dion PA, Bercier V, Bourgouin P, Rochefort D, Bel Hadj S, Durham HD, Velde CV (2010) Gain and loss of function of ALS-related mutations of TARDBP (TDP-43) cause motor deficits in vivo. Human molecular genetics 19 (4):671–683

40. Kabashi E, Bercier V, Lissouba A, Liao M, Brustein E, Rouleau GA, Drapeau P (2011) FUS and TARDBP but not SOD1 interact in genetic models of amyotrophic lateral sclerosis. PLoS genetics 7 (8)

41. Laird AS, Van Hoecke A, De Muynck L, Timmers M, Van Den Bosch L, Van Damme P, Robberecht W (2010) Progranulin is neurotrophic in vivo and protects against a mutant TDP-43 induced axonopathy. PLoS One 5 (10)

42. Ramesh T, Lyon AN, Pineda RH, Wang C, Janssen PM, Canan BD, Burghes AH, Beattie CEJDm, mechanisms (2010) A genetic model of amyotrophic lateral sclerosis in zebrafish displays phenotypic hallmarks of motoneuron disease. 3 (9-10):652–662

43. Robinson KJ, Yuan KC, Don EK, Hogan AL, Winnick CG, Tym MC, Lucas CW, Shahheydari H, Watchon M, Blair IP (2019) Motor Neuron Abnormalities Correlate with Impaired Movement in Zebrafish that Express Mutant Superoxide Dismutase 1. Zebrafish 16 (1):8–14

44. Sakowski SA, Lunn JS, Busta AS, Oh SS, Zamora-Berridi G, Palmer M, Rosenberg AA, Philip SG, Dowling JJ, Feldman ELJMn (2012) Neuromuscular effects of G93A-SOD1 expression in zebrafish. 7 (1):44

45. Schmid B, Hruscha A, Hogl S, Banzhaf-Strathmann J, Strecker K, van der Zee J, Teucke M, Eimer S, Hegermann J, Kittelmann M (2013) Loss of ALS-associated TDP-43 in zebrafish causes muscle degeneration, vascular dysfunction, and reduced motor neuron axon outgrowth. Proceedings of the National Academy of Sciences 110 (13):4986–4991

46. Shaw MP, Higginbottom A, McGown A, Castelli LM, James E, Hautbergue GM, Shaw PJ, Ramesh TM (2018) Stable transgenic C9orf72 zebrafish model key aspects of the ALS/FTD phenotype and reveal novel pathological features. Acta Neuropathologica Communications 6 (1):125

47. Svahn AJ, Don EK, Badrock AP, Cole NJ, Graeber MB, Yerbury JJ, Chung R, Morsch M (2018) Nucleo-cytoplasmic transport of TDP-43 studied in real time: impaired microglia function leads to axonal spreading of TDP-43 in degenerating motor neurons. Acta neuropathologica 136 (3):445–459

48. Tsien RY, Bacskai BJ, Adams SR (1993) FRET for studying intracellular signalling. Trends in cell biology 3 (7):242–245

49. Xu Y, Piston DW, Johnson CH (1999) A bioluminescence resonance energy transfer (BRET) system: application to interacting circadian clock proteins. Proceedings of the National Academy of Sciences 96 (1):151–156

50. Ghosh I, Hamilton AD, Regan L (2000) Antiparallel leucine zipper-directed protein reassembly: application to the green fluorescent protein. Journal of the American Chemical Society 122 (23):5658–5659

51. Kodama Y, Hu C-DJB (2012) Bimolecular fluorescence complementation (BiFC): a 5-year update and future perspectives. 53 (5):285–298

52. Shyu YJ, Hu C-D (2008) Fluorescence complementation: an emerging tool for biological research. Trends in biotechnology 26 (11):622–630

53. Kerppola TK (2006) Design and implementation of bimolecular fluorescence complementation (BiFC) assays for the visualization of protein interactions in living cells. Nature protocols 1 (3):1278

54. Chun W, Waldo GS, Johnson GV (2007) Split GFP complementation assay: a novel approach to quantitatively measure aggregation of tau in situ: effects of GSK3β activation and caspase 3 cleavage. Journal of neurochemistry 103 (6):2529–2539

55. Tak H, Haque MM, Kim MJ, Lee JH, Baik J-H, Kim Y, Kim DJ, Grailhe R, Kim YK (2013) Bimolecular fluorescence complementation; lighting-up tau-tau interaction in living cells. PLoS One 8 (12):e81682–e81682. doi:10.1371/journal.pone.0081682

56. Harvey SA, Smith JCJPB (2009) Visualisation and quantification of morphogen gradient formation in the zebrafish. 7 (5):e1000101

57. Maurya AK, Tan H, Souren M, Wang X, Wittbrodt J, Ingham PWJD (2011) Integration of Hedgehog and BMP signalling by the engrailed2a gene in the zebrafish myotome. 138 (4):755–765

58. Krishnakumar P, Riemer S, Perera R, Lingner T, Goloborodko A, Khalifa H, Bontems F, Kaufholz F, El-Brolosy MA, Dosch RJPg (2018) Functional equivalence of germ plasm organizers. 14 (11):e1007696

59. Moustaqil M, Fontaine F, Overman J, McCann A, Bailey TL, Rudolffi Soto P, Bhumkar A, Giles N, Hunter DJB, Gambin YJNar (2018) Homodimerization regulates an endothelial specific signature of the SOX18 transcription factor. 46 (21):11381–11395

60. Feng S, Varshney A, Villa DC, Modavi C, Kohler J, Farah F, Zhou S, Ali N, Müller JD, Van Hoven MK (2019) Bright split red fluorescent proteins for the visualization of endogenous proteins and synapses. Communications biology 2 (1):1–12

61. Vicario M, Cieri D, Vallese F, Catoni C, Barazzuol L, Berto P, Grinzato A, Barbieri L, Brini M, Calì T (2019) A split-GFP tool reveals differences in the sub-mitochondrial distribution of wt and mutant alpha-synuclein. Cell death & disease 10 (11):1–16

62. Aelvoet S-A, Ibrahimi A, Macchi F, Gijsbers R, Van den Haute C, Debyser Z, Baekelandt Vjjon (2014) Noninvasive bioluminescence imaging of α-synuclein oligomerization in mouse brain using split firefly luciferase reporters. 34 (49):16518–16532

63. Kiechle M, von Einem B, Höfs L, Voehringer P, Grozdanov V, Markx D, Parlato R, Wiesner D, Mayer B, Sakk Ojcr (2019) In Vivo Protein Complementation Demonstrates Presynaptic α-Synuclein Oligomerization and Age-Dependent Accumulation of 8–16-mer Oligomer Species. 29 (9):2862-2874. e2869

64. Kodama Y, Hu C-D (2010) An improved bimolecular fluorescence complementation assay with a high signal-to-noise ratio. Biotechniques 49 (5):793–805

65. Westerfield M (2000) A guide for the laboratory use of zebrafish (Danio rerio). The zebrafish book 4

66. Trinh R, Gurbaxani B, Morrison SL, Seyfzadeh M (2004) Optimization of codon pair use within the (GGGGS) 3 linker sequence results in enhanced protein expression. Molecular immunology 40 (10):717–722

67. Don EK, Formella I, Badrock AP, Hall TE, Morsch M, Hortle E, Hogan A, Chow S, Gwee SS, Stoddart Jjjz (2017) A Tol2 gateway-compatible toolbox for the study of the nervous system and neurodegenerative disease. 14 (1):69–72

68. Morsch M, Radford RA, Don EK, Lee A, Hortle E, Cole NJ, Chung Rsjj (2017) Triggering cell stress and death using conventional UV laser confocal microscopy. (120):e54983

69. Formella I, Svahn AJ, Radford RA, Don EK, Cole NJ, Hogan A, Lee A, Chung RS, Morsch Mjrb (2018) Real-time visualization of oxidative stress-mediated neurodegeneration of individual spinal motor neurons in vivo. 19:226–234

70. Morsch M, Radford R, Lee A, Don EK, Badrock AP, Hall TE, Cole NJ, Chung R (2015) In vivo characterization of microglial engulfment of dying neurons in the zebrafish spinal cord. Frontiers in cellular neuroscience 9:321

71. Tadros W, Lipshitz HD (2009) The maternal-to-zygotic transition: a play in two acts. Development 136 (18):3033–3042

72. Mathavan S, Lee SG, Mak A, Miller LD, Murthy KRK, Govindarajan KR, Tong Y, Wu YL, Lam SH, Yang H (2005) Transcriptome analysis of zebrafish embryogenesis using microarrays. PLoS genetics 1 (2):e29

73. Ayala YM, Zago P, D’Ambrogio A, Xu Y-F, Petrucelli L, Buratti E, Baralle FE (2008) Structural determinants of the cellular localization and shuttling of TDP-43. Journal of cell science 121 (22):3778–3785

74. Chun W, Waldo GS, Johnson GV (2010) Split GFP complementation assay for quantitative measurement of tau aggregation in situ. In: Alzheimer’s Disease and Frontotemporal Dementia. Springer, pp 109–123

75. Foglieni C, Papin S, Salvadè A, Afroz T, Pinton S, Pedrioli G, Ulrich G, Polymenidou M, Paganetti P (2017) Split GFP technologies to structurally characterize and quantify functional biomolecular interactions of FTD-related proteins. Scientific reports 7 (1):14013. doi:10.1038/s41598-017-14459-w

76. George-Hyslop P, Lin JQ, Miyashita A, Phillips E, Qamar S, Randle S, Wang G (2018) The Physiological and Pathological Biophysics of Phase Separation and Gelation of RNA Binding Proteins in Amyotrophic Lateral Sclerosis and Fronto-Temporal Lobar Degeneration. Brain Research 1693. doi:10.1016/j.brainres.2018.04.036

77. Matsumoto T, Matsukawa K, Watanabe N, Kishino Y, Kunugi H, Ihara R, Wakabayashi T, Hashimoto T, Iwatsubo T (2018) Self-assembly of FUS through its low-complexity domain contributes to neurodegeneration. Human molecular genetics 27 (8):1353–1365. doi:10.1093/hmg/ddy046

78. Vance C, Scotter EL, Nishimura AL, Troakes C, Mitchell JC, Kathe C, Urwin H, Manser C, Miller CC, Hortobágyi T, Dragunow M, Rogelj B, Shaw CE (2013) ALS mutant FUS disrupts nuclear localization and sequesters wild-type FUS within cytoplasmic stress granules. Human molecular genetics 22 (13):2676–2688. doi:10.1093/hmg/ddt117

79. Mutihac R, Alegre-Abarrategui J, Gordon D, Farrimond L, Yamasaki-Mann M, Talbot K, Wade-Martins Rjnod (2015) TARDBP pathogenic mutations increase cytoplasmic translocation of TDP-43 and cause reduction of endoplasmic reticulum Ca2+ signaling in motor neurons. 75:64–77

80. Masuda-Suzukake M, Nonaka T, Hosokawa M, Kubo M, Shimozawa A, Akiyama H, Hasegawa M (2014) Pathological alpha-synuclein propagates through neural networks. Acta Neuropathologica Communications 2 (1):88. doi:10.1186/s40478-014-0088-8

81. Alberti S, Gladfelter A, Mittag TJC (2019) Considerations and challenges in studying liquid-liquid phase separation and biomolecular condensates. 176 (3):419–434

82. Elbaum-Garfinkle Sjjobc (2019) Matter over mind: Liquid phase separation and neurodegeneration. 294 (18):7160–7168

