## supplemental data for "*In Vivo* Validation of Bimolecular Fluorescence Complementation (BiFC) to Investigate Aggregate Formation in Amyotrophic Lateral Sclerosis (ALS)"

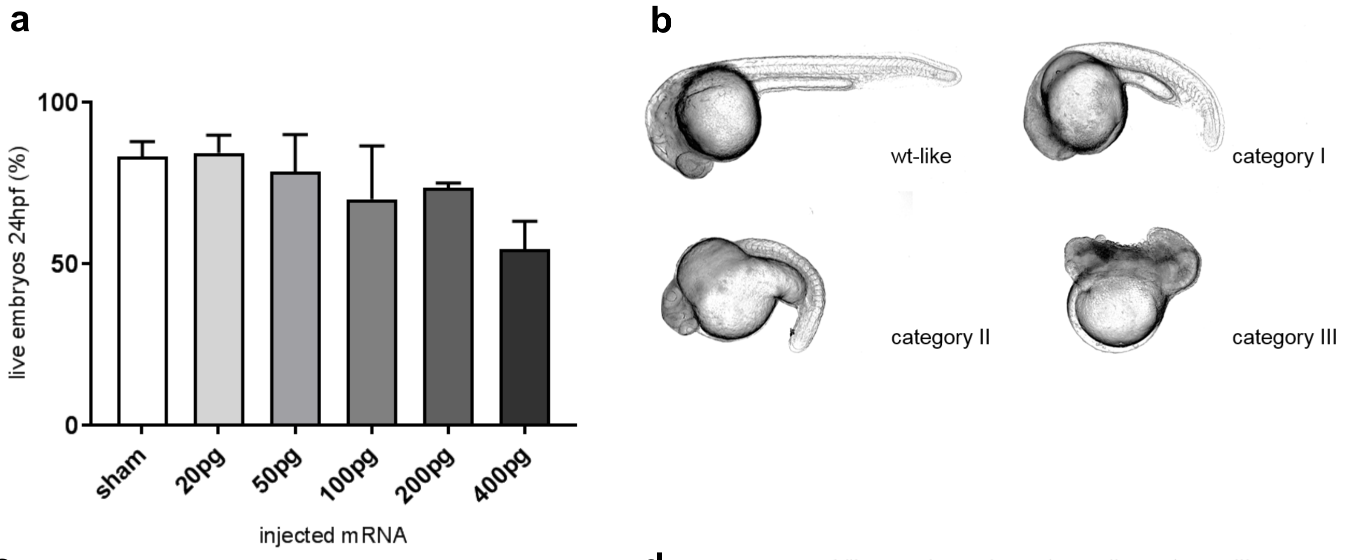


c

| | |
|---|---|
| **wt-like** | morphology not distinguishable from non-injected larvae |
| **category I** | minor malformations *(slight spinal curvature and/or modestly decreased head size)* |
| **category II** | prominent malformations *(strong spinal curvature, yolk sack deformation, underdeveloped head & eyes)* |
| **category III** | severe malformations *(shortened body, poorly developed head & tail, opaque tissue)* |

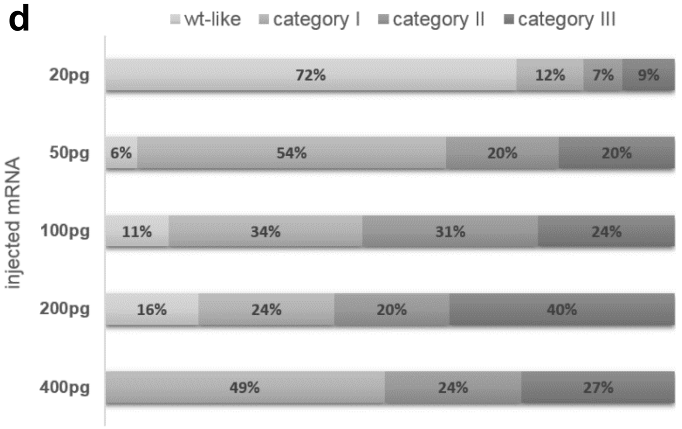


**Supplemental Fig 1 Effect of TDP-43 BiFC mRNA microinjections on zebrafish survival and morphology**

**a**: Zygotes at one-cell stage were microinjected with different amounts of wtTDP-43 BiFC mRNA and screened for survival at 24 hpf. Doses refer to the total of both complementary mRNAs. Results are shown as percentage of non-injected control embryos, and data are pooled from 3 independent experiments. **b**: Representative pictures and classification of morphological abnormalities observed at 24 hpf after wtTDP-43 BiFC mRNA microinjections. **c**: Description of morphological criteria used to categorize wtTDP-43 BiFC mRNA microinjected embryos. **d**: Quantitative evaluation of morphological phenomena in wtTDP-43 BiFC mRNA microinjected embryos at 24hpf, based on data from 3 independent experiments with a minimum of n=20 for each dose.

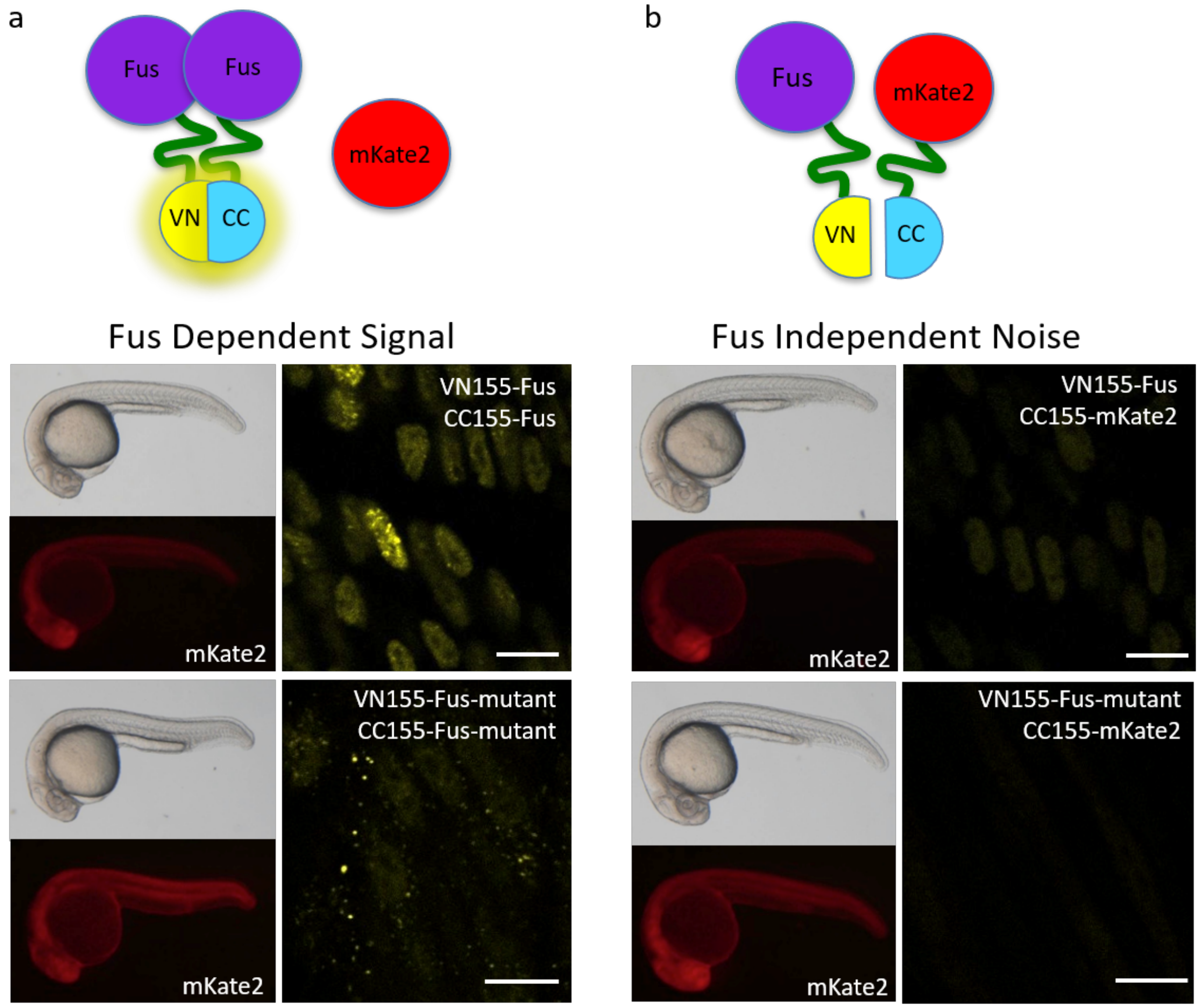


**Supplemental Fig 2 Fus aggregation in zebrafish is specific**

Fus-BiFC complementation assays. **a:** Fus dependent signal: 400 pg H2B-mCerulean3, 100 pg VN155-Fus (VN155-Fus-mutant), 100 pg CC155-Fus (CC155-Fus-mutant) and 40 pg mKate2 mRNA were co-injected into 1-cell stage wildtype embryos. Representative pictures of the zebrafish wild-type Fus (top panel) and mutant Fus (bottom panel) BiFC signal at 28 hpf in the somites over the yolk extension. Nuclear or cytoplasmic Fus-mVenus signal is detected in Fus dependent assays. High-magnification images also used in Figure 4. **b:** Fus independent noise: 400 pg H2B-mCerulean3, 100 pg of VN155-Fus (or VN155-Fus-mutant) and 50 pg of CC155-mKate2 mRNA were co- injected into 1-cell stage wildtype embryos. Representative pictures of the zebrafish wild-type Fus/mKate2 (top panel) and mutant Fus/mKate2 (bottom panel) BiFC signal at 28 hpf in the somites over the yolk extension. Scale bars represent 10 µm.
